## Supplementary material for "Bacteria face trade-offs in the decomposition of complex biopolymers": S1 Text

### I. METHODS

#### A. Prevalence of different chitinase classes in bacteria

##### 1. *Screening of chitinase prevalence in genomes of bacteria*

To investigate the genomic potential of bacteria to produce/house/have chitinases (Fig A), with given KEGG identifiers, we screened the genomes for selected enzymes via two tools: the first one for targeting soil bacteria was JGI IMG Integrated Microbial Genomes & Microbiomes (IMG/M) tool (https://img.jgi.doe.gov/) accessed on 19.01.2023; and a second for a broader search targeting all environments was Anotree (http://annotree.uwaterloo.ca/) accessed on 11.11.2022.

For JGI, all available bacterial genomes ('Isolates') were subset by selecting those listed as 'Soil' in the category 'Ecosystem Type' resulting in 4034 bacterial genomes. These genomes were imported into the genome cart and screened for the presence of KEGG functions under 'Find Functions'. Here, the respective KEGG number (e.g. K01207) was added under 'Keyword' and 'KEGG Orthology ID (list)*' under 'Filters'. The KEGG function, name, definition, and resulting genome hits are listed in Table A. Since some of the hits correspond to different strains of the same species we have grouped them, and assumed a single count for such groups for the analysis in the main text. We proceeded in an analogous way using the Anotree, there screening the whole database. As a result we obtained a list of all species that have the potential to produce each one of the chitin degrading enzymes. For the main analysis we selected only some identifiers highlighting them in Table A.

| KO ID | KO Name | KO Definition | Genome hits Anotree |
|---|---|---|---|
| K01183 | E3.2.1.14 | chitinase [EC:3.2.1.14] | 5189 |
| K01207 | nagZ | beta-N-acetylhexosaminidase [EC:3.2.1.52] | 18609 |
| K01233 | csn | chitosanase [EC:3.2.1.132] | 934 |
| K01452 | E3.5.1.41 | chitin deacetylase [EC:3.5.1.41] | 7079 |
| K03791 | - | putative chitinase | 2277 |
| K15855 | csxA | exo-1,4-beta-D-glucosaminidase [EC:3.2.1.165] | 954 |

TABLE A: List of enzymes involved in chitin degradation, their identifiers and the genome hits in the two databases. We highlight (in cyan) the chitinases analyzed in the main text in Fig 2.

##### 2. *Literature Search: Experimental prevalence of exo/endo-chitinases in bacteria*

A systematic literature search was performed to assess the experimentally tested prevalence of exo- and endo-chitinases in microorganisms. The search term "Chitinase Assay Kit" was used in the literature search engine "Google

*
†

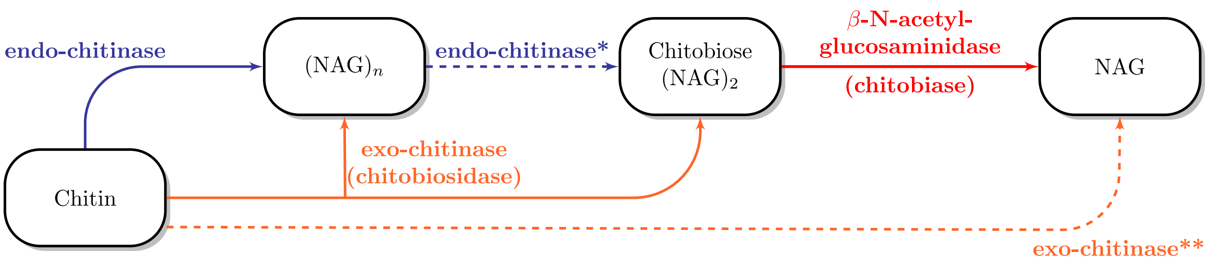


FIG. A: Schematic representation of the hydrolysis of chitinase into NAG (N-acetylglucosamine), which involves: endo- and exo-chitinases, as well as a N-acetylglucosaminidase. Note that to maintain the broad scope of our research, we do not incorporate all the specific sets of reactions into our degradation model. [a] [b]

[a] Endo-chitinases are also able to cut dimers from the chain however this reaction may have a different turnover rate.
[b] There is a potential for certain types of exo-chitinases to cleave directly monomers from the long chains of polymers.

Scholar" (https://scholar.google.com/) to identify studies that test the ability of different microorganisms to specifically degrade chitobiose (to detect exochitinase production) or chitotriose (to detect endochitinase production). Studies that tested the chitinase production of fungi were not included, as our study mainly focuses on bacteria. All selected studies either used p-Nitrophenol- or 4-Methylumbelliferone (4-MUF)-marked oligomers to quantify exo- or endochitinase activity colormetrically or fluormetrically. We then constructed a table containing the information on the species and strains used in each study, and whether they observed exo- or endo-chitinase activity in a binary way ("1" for activity and "0" for no activity). Whenever chitinase activity was given in a quantitative way, a threshold was applied to the respective value because the values often spanned multiple orders of magnitude. For example, in the study by Bittleston et al. (2020) every value above 5U was chosen as the threshold above which we assigned the presence of exo- or endochitinase activity ("1"). This threshold was evaluated for each study individually, as they often differed in the used units and experimental approaches. The resulting table was then used to create Fig 2C of the main text, by assigning groups ("exo" when only exo-chitinase activity was observed, "endo" when only endo-chitinase activity was observed and "both" when both types of enzymes were observed) and aggregating the data on species level, calculating the relative abundance of each group for each species across all studies where it occurred. A list of the used studies[1–4, 6–8, 11, 14, 15, 18–22, 24–28] is provided in supplementary file S1_Overview.csv.

### B. Model description

The general elements of the model are: microorganisms, enzymes, and nutrients (in the form of polymers of different sizes and dissolved inorganic nitrogen). In the next subsections we describe some of its main elements. The full list of all the parameters their meaning and units in Table D.

**Kinetics of exo-enzymes.** Considering $N_i$ the concentration of a i-mer, and the $E_{exo}$ the concentration of exo-chitinase in solution, the size distribution of polymers of maximum size $n$ changes as:

$$N_n(t+dt) = N_n(t) - N_n f(E_{exo})dt \tag{1}$$

$$N_i(t+dt) = N_i(t) - N_i(t)f(E_{exo})dt + N_{i+1}(t)f(E_{exo})dt, \quad \text{for} \quad (2 < i < n) \tag{2}$$

$$N_1(t+dt) = N_1(t) + 2N_2(t)f(E_{exo})dt + \sum_{i=3}^{N} N_i(t)f(E_{exo})dt, \tag{3}$$

where

$$f(E_{exo}, t) = \frac{k_{exo} \cdot E_{exo}(t)}{k_m + \sum_{i=1}^{n} N_i(t)}, \tag{4}$$

where $k_m$ and $k_{exo}$ are respectively the half saturation constant and the turnover number of exo-chitinase. Note that the reaction kinetics $f(E_{exo})$ is the same for all molecules in the solution, and is independent of the molecule sizes. Also note that the reaction depends on the total number of polymers in solution, and the enzymes kinetics saturates with the total number of molecules in solution, the total concentration $\sum_{i=1}^{n} N_i$ given in fmol C/mm$^3$.

**Kinetics of endo-enzymes.** The evolution of the size distribution of polysaccharides under the action of the endo-enzymes is modeled by:

$$N_i(t+dt) = N_i(t) - N_i(t) f_i'(E_{endo}, t)dt + \sum_{j>i}^{n-1} g(i,j) f_j'(E_{endo}, t) N_j(t)dt, \tag{5}$$

where $g(i,j)$ represents the fragment distribution function, i.e. it gives the probability of molecules of size i produced by the hydrolysis of a molecule of size j. Given the binary nature of the reaction this function is defined by:

$$g(i,j) = \frac{2}{j-1}. \tag{6}$$

Additionally the reaction kinetics can be described by:

$$f_j'(E_{endo}, t) = \frac{(j-1)k_{endo} \cdot E_{endo}(t)}{k_m' + \sum_{k=2}^{n}(k-1)N_k(t)}, \tag{7}$$

note that here in contrast to the previous model, the $f_j'(E_{endo})$ depends on the size of the molecule, since larger molecules have more bonds which can be attacked by endo-enzymes. The factor $j-1$ in the numerator refers to the number of forms a polymer of size $j$ can be broken (e.g. $j=5$, can be broken in 4 ways: 2 and 3 ($\times 2$); 1 and 4 ($\times 2$)). The factor $k-1$ in the denominator sum, however refers to all possible bonds that can be attacked by an endo-enzyme. All parameters used for enzyme kinetics are listed in Table D.

**Microbial growth dynamics**. The model consists of various spatially and temporally defined elements: microbial biomass $C_b$ (in mol C/mm$^3$), available enzyme $E_i$ (where $i$ defines either *exo* or *endo* enzyme type), available substrate $N_i$( where $i$ defines the polymer size) and finally the amount of dissolved inorganic nitrogen DIN. The elements in each microsite evolve to account for enzyme and microbial activity. Each model component is defined by its C:N ratio, which we list in Tab. B.

| component of the model | parameter | C:N ratio |
|---|---|---|
| microorganisms | $\mathrm{CN}_B$ | 11.3 |
| enzyme | $\mathrm{CN}_E$ | 5 |
| substrate (polymers and monomers) | $\mathrm{CN}_S$ | 8 |

TABLE B: The C:N ratios used in simulations.

Microorganisms in particular are considered to be constituted by three major fractions: protein rich fraction with $\mathrm{CN}_P$ ($f_1$); carbon rich fraction characterized by $\mathrm{CN}_C$ ($f_2$) and a small monomer fraction with $\mathrm{CN}_S$ ($f_3$). The microbial C:N ratio is set as:

$$\mathrm{CN}_B = \frac{\text{Number of C molecule}}{\text{Number of N molecules}} = \frac{\sum_{i=1}^{3} f_i}{\sum_{i=1}^{3} \frac{f_i}{\mathrm{CN}_i}} = \frac{1}{\sum_{i=1}^{3} \frac{f_i}{\mathrm{CN}_i}}. \tag{8}$$

After microbial death these fractions are re-distributed into different pools of substrates, see Fig B.

In each time steps microorganisms: (i) uptake monomers $N_1(t)$, (ii) keep their general function (maintenance), (iii) produce enzymes, (iv) grow, (v) die and (vi) reproduce. We follow with a detail description of each step and all parameters involved.

**(i) Uptake.** The uptake dynamics is slightly modified from [13], therefore we provide more details on it. We assume that microorganisms take up only monomers. The uptake requires membrane transporters and the uptake rate depends on both the concentration of monomers and the number of transporters on the cell membrane. In the limit case with very high concentration of substrate (monomers) the amount transported in a time interval will be proportional to the surface area of the cell. The maximum possible uptake for a single cell is calculated according to:

$$U_{cell} = u \cdot f_r \cdot R_{sv} \cdot V \quad [\text{mol C h}^{-1}\ \text{cell}^{-1}] \tag{9}$$

where $V$ is the volume of one microbial cell, $R_{sv}$ is the surface-volume ratio, $f_r$ is the fraction of cell membrane occupied by channels, $u$ is the conductivity of a single channel. The parameter $u$ can be controlled in the simulation. We assume that a bacterial cell has 10% of its area covered by channels, i.e $f_r = 0.1$ [5, 9]. The surface-volume ratio and the cell volume change as the cell size grows. Knowing the average cell density, it is easy to estimate the total volume of cells from the cell's biomass, $V = \frac{\text{cell biomass}}{\text{cell density}}$. Note that the model does not track the total biomass but only the carbon fraction $f_c$, which accounts only for 10% of the cell biomass [23]. Given that the molecular weight of carbon is 12 fg/fmol we can compute the total biomass from the cell's carbon b:

$$\text{biomass} = \frac{b \cdot 12}{f_c} = \frac{b \cdot 12}{0.1} = 1.2 \cdot b \quad [\text{fg}] \tag{10}$$

Taking into account the density of the cell $\rho = 1000$ fg/$\mu$m$^3$ [17] and Eq.(10), we can compute the volume. The surface to volume ratio can be easily computed by assuming a spherical cell $R_{sv} = \frac{3}{\left(\frac{3V}{4\pi}\right)^{1/3}}$. Taking into account the information above the equation for $U_{cell}$ of a cell as a function of stored carbon is given by:

$$U_{cell} = u \cdot k_1 \cdot b^{2/3} 8 \quad [\text{mol C h}^{-1}\text{cell}^{-1}], \tag{11}$$

where $k_1 = 0.00546$, considering the biomass of a single cell $b$ and considering the uptake of chitin monomers (NAG) with 8 carbon atoms. Considering that the microsite contains multiple cells $n_{\text{col}}$, and with a biomass of $C_b$. We estimate the number of cells within the microsite as $n_{\text{col}} = C_b/b$, where $b$ defines the average biomass of a single cell, which we consider to be approximately 4 fmol C. The uptake of the whole colony would be defined as $U'_{pot} = U_{cell} n_{\text{col}}$, or:

$$U'_{pot} = u \cdot k_1 \cdot C_b^{2/3} 8 \cdot n^{1/3} \quad [\text{mol C h}^{-1}]. \tag{12}$$

In the case of a limited substrate the uptake in a time step for the microbes inhabiting one microsite depends both on the substrate and receptor numbers, and we assume Monod like kinetics:

$$U_C = \frac{U'_{pot} N_1(t)}{k_{up} + N_1(t)}, \tag{13}$$

where $N_1(t)$ represent the concentration of monomers in the environment at time $t$. This value gives the mol C of substrate so we directly set up the amount of carbon taken up as $U_C$ in mol C, and the amount of nitrogen as $U_N = U_C/$ CN$_S$ mol N. All symbols used for microbial model are listed in Tab. D.

**(ii) Maintenance.** As in [12, 13] maintenance only requires carbon, this carbon is linearly proportional to the biomass $r_m C_b$. If the uptake has not enough carbon to meet maintenance requirements, the necessary carbon is subtracted from biomass.

**(iii) Enzyme production.** After maintenance, we follow with enzyme production. Microorganisms use a fraction $e_{fr}$ of the taken up carbon for enzyme production, to that we also add an associated respiration cost to produce enzymes $r_e$. Given the estimation that up to 75% of bacterial energy is used for protein production [16], we set up the $efr = 0.5$. Therefore the total carbon invested to produce $e_{fr}(U_C - r_m C_b)$ of enzymes is $e_{fr}(U_C - r_m C_b)/(1 - r_e)$. Note that for our parameter choice (i.e. for any $(1 - r_e) > e_{fr}$) enzyme production takes only a fraction of the available carbon (what is left from the uptake). The other necessary condition for the production of enzyme is the availability of nitrogen. If the enzyme production is N-limited, enzymes are only produced to the limit $U_N \cdot \text{CN}_E$. In other words, the amount of enzymes produced $\Delta E_i$ at each time step is given by:

$$\Delta E_i = \begin{cases} \min\left[e_{fr}(U_C - r_m C_b), U_N \cdot \text{CN}_E\right] & \text{if} \quad U_C - r_m C_b > 0, \\ 0 \quad \text{else}, \end{cases} \tag{14}$$

where $i$ can be *exo* or *endo*, or in the case of simultaneous production it represents the total of enzymes produced$\Delta E_T$. This total amount is divided in the two fractions $fr \cdot \Delta E_T = \Delta E_{exo}$ and $(1 - fr) \cdot \Delta E_T = \Delta E_{endo}$. In this work we assume that enzyme production fully depends on the uptake, and enzymes cannot be produced at the costs of biomass.

**(iv) Growth.** Finally if there is N and C left after enzyme production the rest goes into growth. In other words we need $U_C - r_m C_b - \frac{\Delta E_i}{(1-r_e)} > 0$ and $U_N - \Delta E_i/CN_E > 0$ to be able to grow. The possible growth is established by the limiting resource C or N. Again, in an analogous way to enzyme production, there is a respiration association with growth $r_g$. All parameters and their corresponding values are listed in Tab. D.

$$\Delta C_b = \min\left[\left(U_C - r_m C_b - \frac{\Delta E_i}{(1 - r_e)}\right)(1 - r_g), U_N CN_E - \Delta E_i\right] \quad \text{if} \quad U_C - r_m C_b - \frac{\Delta E_i}{(1 - r_e)} > 0 \tag{15}$$

In case there are no more resources left, there is no growth and $\Delta C_b = 0$.

$$C_b(t + dt) = C_b(t) + \Delta C_b. \tag{16}$$

**(v) Death of microorganisms.** We take into account stochastic death events, which constitute of a random elimination of 1% of microbial biomass per hour. Another reason for death can be starvation, if microbial biomass reaches below a predetermined limit ($B_{min}$, values in Tab. D) microorganisms cannot survive further. After death microbial biomass is divided in the corresponding C-rich, N-rich and monomer-rich pools, see Fig B. Only the monomer pool can be reused by other microorganisms around it.

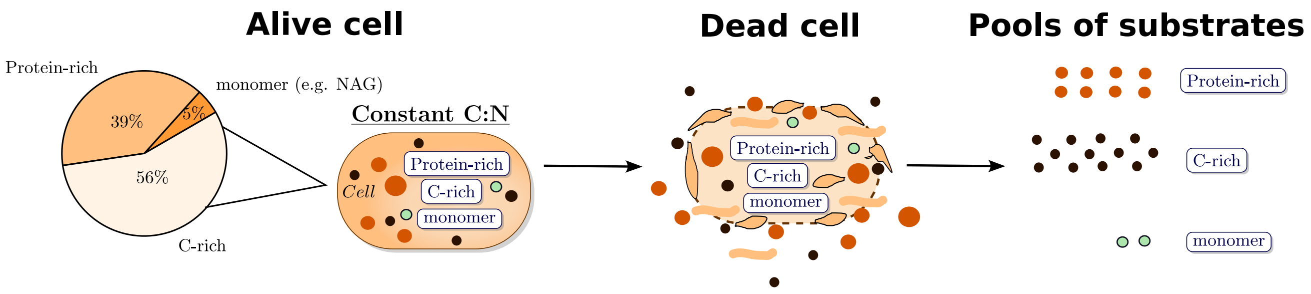


FIG. B: **Overview of re-distribution of microbial remains into different substrate pools after microbial death.** Microorganisms keep their C:N ratio constant (11.3), we consider that they consist mainly of protein-rich (C:N ratio 5), C-rich components (C:N ratio 150) and few monomers (C:N 8). When microorganisms die these pools are released into the environment together with small amounts of monomers. To avoid mixing the effect of degradation of necromass with depolymerization on growth we assume that microorganisms in the system cannot survive on microbial remains. The enzymes produced also have a fixed C:N ratio of 5, after decay these enzymes also enter the protein-rich fraction of substrate pool, but cannot be recycled by the current type of microorganisms. Finally the polymers present in the environment have C:N ratio 8, which does not change with fragmentation dynamics.

**(vi) Reproduction.** As described in the main text the system is constituted by a spatial grid ($100 \times 100$) microsites. Microorganisms are considered of limited motility, they only spread in space to neighboring microsites when population becomes large. We assume that a small colony of microorganisms can occupy one microsite. We assume that bacterial cells of about 500 fg [10]. Considering that 10% of cell is composed by carbon and the carbon molecular mass of 12 fg/fmol a cell can be assumed to be on average around $b = 4$ mol C. A microsite of $10^3$ $\mu$m$^3$, may contain approximately $10^3$ densely packed cells, however we assume that the volume is divided among the colony and the other components of the environment (soil matrix and organic matter). We assume that a microsite-colony can contain at most the biomass of 3 average-size cells (Bmax = 3) without attempting to spread in space. Once the biomass of a microsite exceeds 12 mol C/MS (indicating the presence of 4 cells or more), the colony would divide. Note that the algorithm does not track how the biomass is exactly divided within the members of a single micro-site colony, and the number of members within this colony is only an estimation based on the average-cell size provided. After a limit of biomass is exceeded, a new directly neighboring microsite is selected by random and colonized. In this step the biomass is divided in half, where the first half remains in the current site and the other half is transferred to the neighboring microsite. In case there are no free microsites available the biomass continues to grow, filling the left space, and divides at the next available opportunity.

**Enzyme decay.** Finally, extracellular enzymes also become inactive after some time with the rate $k_e$. The change of enzymes in a time step is given by

$$E_i(t + dt) = E_i(t) - k_e E_i(t) + \Delta E_i. \tag{17}$$

**Spatial dynamics** Large polymers are insoluble and therefore cannot diffuse. On the other hand oligomers with sizes smaller than 10 units do diffuse, but their diffusion coefficient decreases with size, as represented in Fig C. Finally we also take into account the diffusion of dissolved inorganic nitrogen DIN.

#### C. Simulation dynamics and analysis

All microsites are initialized with the same amounts of polymers, all of which have the same size $n$. We start with few microorganisms in the central region of the grid. Each microorganism also starts with an initial amount of enzymes, which is necessary to give an initial boost to the decomposition process. It is important to note that there are different ways to trigger the initial growth, which could be the direct availability of a certain amount of monomers

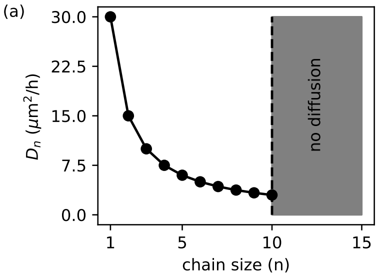


FIG. C: The diffusion rate of oligomers decreases with their size, $n$. Note that the scaling down of $D_n$ accounts only for how the coefficient changes with molecular weight of the polymers, however the shape as well as the porosity of the soil matrix may further slow down the diffusion rates.

or as we did with the presence of enzymes. Although the quantitative aspects of the simulation results depend on initial conditions, the qualitative facets do not.

The temporal dynamics is advanced by a finite difference method. Finite differences are also used to implement spatial dynamics to model the diffusion process. The step size was adapted to guarantee the stability of both the diffusion and fragmentation processes, since numerical instabilities can lead to violation of conservation of mass in the system. We have chosen the values for dt from 0.05 h to 0.005 h depending on the set-up. The smallest dt values were used to simulation the dynamics with high diffusion coefficients.

| variable | symbol | initial value well mixed | initial value spatial model | units |
|---|---|---|---|---|
| Microbial biomass/microsite | $C_b(0)$ | 4 | 1 | nmol C/ mm$^3$ |
| Enzymes/microsite | $E_i(0)$ | 1 | 1(microsite with microbe)<br>0.001 (empty microsite) | nmol C/mm$^3$ |
| Polymers/microsite | $N_n(0)$ | – | – | nmol/ mm$^3$ |
| Monomers | $N_1(0)$ | 0 | 0 | nmol/ mm$^3$ |
| DIN | DIN(0) | 0.5 | 0.5 | nmol/ mm$^3$ |

TABLE C: Initial concentrations for all the components of the model in the well mixed and spatially explicit scenarios.

### II. ADDITIONAL RESULTS

**Degradation dynamics** In this part we present some additional results of the comparison of degradation dynamics of endo and exo-enzymes. Here we compare the necessary time to degrade different pools of complex substrate. We assume that all pools have exactly the same amount of nutrients $M = 200$ nmol /mm$^3$ (total amount of monomers), however in each pool all these monomers are trapped within polymeric chains of different lengths. For example in two limit cases we have: either many very short chains; or few very long ones. The amount of nutrients in these cases is the same, but the time necessary to degrade these pools is different. For exo-enzymes the $T_{deg}$ increases linearly with the increase of polymer size in the initial pool, see Fig D. The process takes longer for a single long chain than for many short ones. This occurs because exo-enzymes need free molecule ends to act. Endo-enzymes on the other hand can attack the molecule at any point and for this reason the degradation time $T_{deg}$ is almost independent on the chain size, see Fig D.

**Microbial growth of specialists** Here we present the time series of the microbial growth for high and low nutrient concentrations (150 and 1500 nmol/mm$^3$ respectively). In different experimental setups, microorganisms exhibit either growth or a decline in biomass right from the start. When growth is feasible, microbial biomass undergoes an initial increase up to a maximum point, which subsequently diminishes as the available resources decrease. Notice that

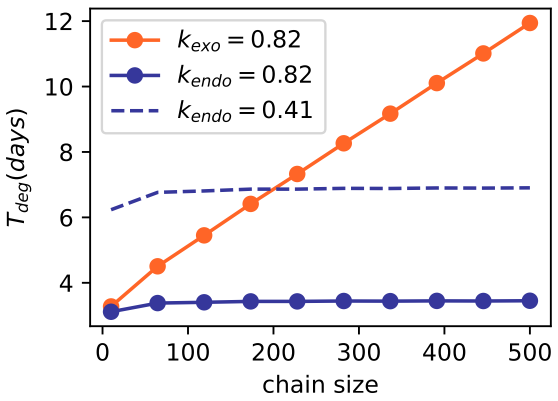


FIG. D: Time to degrade ($T_{deg}$) a pool of substrate consistent of polymers of different chain lengths. The total amount of monomers in the system is fixed to $M = 200$ nmol /mm$^3$, note that as we change the size of the chains present in the initial pool, the number of chains in this pool also changes. The turnover numbers $k_{exo}$ and $k_{endo}$ are given in nmol nmol C$^{-1}$ h$^{-1}$.

the length of polymer chains exerts a more pronounced influence on the growth of exo-producers compared to endo-producers, as demonstrated in Fig E. In low nutrients exo-producers are able to grow only if polysaccharides of small chain length are present. Conversely, under high-nutrient conditions, exo-producers exhibit consistent growth across all chain lengths. Endo-producers, as also explained in the main text only grow in low nutrients, and their biomass decreases when a lot of nutrients is provided.

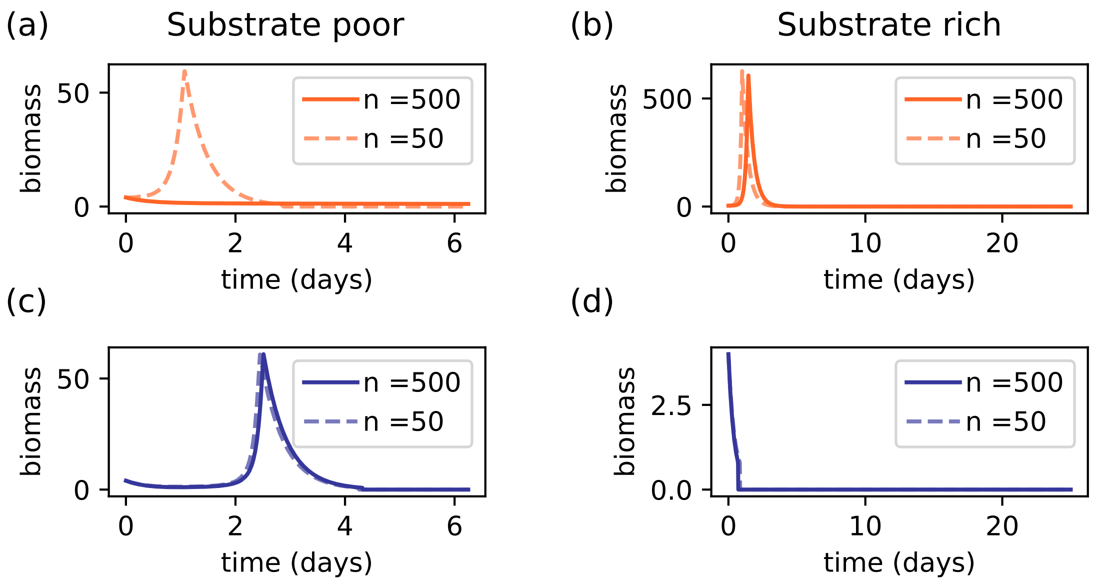


FIG. E: Time series of the biomass of exo producers (in orange) and endo producers (in blue) growting on a pool of polymers of length $n$ (see labels) of low and high nutrient concentrations (with total number of monomers $M = 150$ and $M = 1500$ nmol /mm$^3$ respectively). We consider two scenarios: where these monomers are trapped in chains of small length $n = 50$ with the dynamics represented by the dashed line; and a second scenario where these monomers are trapped in long chains of length $n = 500$ with the dynamics shown by soplid line. Note that the amount of nutrients is the same in the two scenarios, but we have a different total number of chains in the system.

**Microbial growth of generalists** In this section we complement the results of the main text on the growth potential of generalists in different substrate concentrations. Here we present the maximum biomass reached by microorganisms producing $fr_{exo}$ of exo-enzymes, and $1 - fr_{exo}$. Considering that both enzyme have the same turnover number. The results are similar to the ones presented in Fig 6 of the main text.

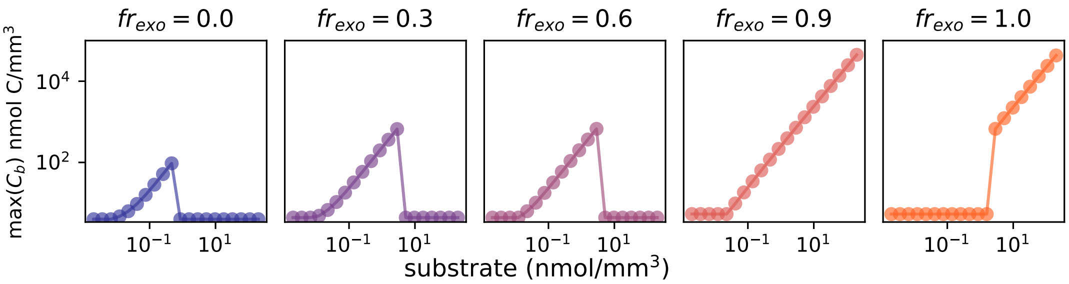


FIG. F: Maximum biomass reached by microorganisms growing on a defined pools of complex substrates with differnet concentrations. Here we consider the same turnover number for endo and exo enzymes ($k_{exo} = k_{endo} = 0.82$). The substrate pools consist of polysaccharides with length $n = 500$.

| Symbols | representation | used value | units |
|---|---|---|---|
| $N_i$ | polymer concentration of size $i$ within a microsite | | nmol C/mm$^3$ |
| $E_i$ | enzyme of type $i$ ($i$ can be *exo* or *endo*) | | nmol C/mm$^3$ |
| $C_b$ | biomass within a microsite | | nmol C/mm$^3$ |
| $k_m$ | half saturation of exo-enzymes | 1. | nmol /mm$^3$ |
| $k'_m$ | half saturation of endo-enzymes | 1. | nmol /mm$^3$ |
| $k_{exo}$ | turnover number of substrate molecules per exo-enzyme | 0.82 | nmol nmol C$^{-1}$ h$^{-1}$ |
| $k_{endo}$ | turnover number of substrate molecules per endo-enzyme | 0.82 or 0.42 | nmol nmol C$^{-1}$ h$^{-1}$ |
| $fr$ | fraction of cell membrane occupied by channels | 0.1 | – |
| Rsv | surface to volume ratio of a cell | | $\mu$m$^{-1}$ |
| V | volume of the cell | | $\mu$m$^3$ |
| $fc$ | carbon fraction | 0.1 | – |
| $\rho$ | density of the cell | 1000 | fg/$\mu$m$^3$ |
| $k_1$ | conversion parameter | 0.00546 | $m^2$ fmol C$^{-2/3}$ |
| $U_{cell}$ | uptake rate possible for a single cell | | mol C h$^{-1}$ cell$^{-1}$ |
| $U'_{pot}$ | uptake rate possible within a microsite | | mol C/h |
| $U_C$ | carbon taken up realized by a microsite-colony, given by Eq.(13) | | mol C/h |
| $U_N$ | nitrogen taken up realized by a microsite-colony | | mol N/h |
| $n_{col}$ | estimation of the cells within the colony | | mol C/h |
| $u$ | conductivity of membrane channels | 1.5 | nmol h$^{-1}$ $\mu$m$^{-2}$ |
| $k_{up}$ | uptake | 1 | nmol/mm$^3$ |
| $r_e$ | resp. enzyme production | 0.05 | h$^{-1}$ |
| $r_g$ | growth resp. | 0.05 | h$^{-1}$ |
| $r_m$ | maintenance resp. | 0.1 | h$^{-1}$ |
| $e_{fr}$ | fraction of C used for enzyme production | 0.5 | - |
| $b$ | size of bacterial cell | 4 | fmol C |
| $B_{max}$ | maximum biomass per microsite | 3 | cells |
| $B_{min}$ | minimum biomass per microsite | 0.02 | cell |
| $d$ | frac. random mortality | 0.01 | of cells h$^{-1}$ |
| $D_0$ | diffusion rate | 30 | $\mu$m$^2$/h |

TABLE D: List of symbols used in the model with the corresponding values used in simulations.
